# Computational assessment of arrhythmia potential in the heterogeneously perfused ventricle

**DOI:** 10.1101/301614

**Authors:** Sanjay R Kharche, Christopher W McIntyre

## Abstract

**Background:** The heterogeneity in the human left ventricle is augmented by heterogeneous perfusion defects in dialysis patients. We hypothesized that ischemic zones generated by heterogeneous perfusion are a cause of clinically observed post-dialysis arrhythmia. This preliminary study assessed the arrhythmia potential in a heterogeneously perfused 2D human ventricle computational model.

**Aim:** Our aim was to ascertain a relationship between the number of ischemia zones and incidence of multiple re-entrant waves in a 2D model of the human ventricle.

**Methods:** A human ventricle action potential model was modified to include the adenosine triphosphate (ATP) sensitive potassium current. Within ischemic zones, cell electrophysiological alterations due to ischemia were implemented as increased extracellular potassium, reduced intracellular ATP concentrations, as well as reduced upstroke current conductances. The cell model was incorporated into a spatial 2D model. The inter-cellular gap junction coupling was adjusted to simulate slow conduction in dialysis patient hearts. CT imaging data of the heart obtained during dialysis was analysed to estimate the approximate spatial size of ischemic zones. An ischemic border zone between the normal and central ischemic zones was implemented which had smoothly varying electrophysiological parameters. Arrhythmic potential was assessed using the paths of the centres of the re-entrant waves, called tip trajectories, and dominant frequency maps.

**Results:** Extracellular potassium elevated the resting potential and I_KATP_ reduced the action potential’s duration. In the absence of ischemic zones, the propensity of the model to induce multiple re-entrant waves was low. The inclusion of ischemic zones provided the substrate for initiation of re-entrant wave fibrillation. The dominant frequency which measured the highest rate of pacing in the tissue increased drastically with the inclusion of ischemic zones, going from 3 Hz in the pre-dialysis state to over 6 Hz in the post-dialysis state. Re-entrant wave tip numbers increased from 1 tip in the pre-dialysis case to 34 in the post-dialysis case, a 34 fold increase. The increase of tip number was found to be strongly correlated to tissue heterogeneity in terms of ischemic zone numbers. Computational factors limiting a more extensive simulation of cause-effect combinations were identified.

**Conclusions:** A dialysis session restores systemic homeostasis, but promotes deleterious arrhythmias. Structure-function mechanistic modelling will permit patient-specific assessment of health status. Such an effort is expected to lead to application of wider physical sciences methods in the improvement of the lives of critically ill patients. High performance computing is a crucial requirement for such mechanistic assessment of health status.

## Introduction

Ventricular arrhythmias are unacceptable clinical complications in patients receiving dialysis. The prevalence of dialysis induced arrhythmias can be as high as 75% [1]. They mostly occur after dialysis as observed by Burton et al. [2]. Whereas dialysis restores systemic serum content, the progressive small vessel disease in the patient’s heart may aggravate ventricular electrical heterogeneity. Clinical therapies for dialysis induce cardiac malfunction as shown by Crowley and McIntyre [3] are continually being developed. However, the structure-function mechanisms of post-dialysis arrhythmias remain unclear, which limits rapid translation of emergent knowledge into clinical practice. It is thought that the arrhythmias may be a manifestation of regionally heterogeneous blood flow, which induces heterogeneous conduction in the myocardium [4]. Heterogeneous conduction has been shown to promote multiple re-entrant waves, also known as spiral waves [5, 6], which are characteristic of arrhythmia. Whereas the genesis and persistence of arrhythmia due to an ischemic zone is known [7, 8], the relationship between multiple ischemic zones of clinically observed dimensions remains underexplored. We hypothesized that the heterogeneous conduction is due to complex ischemic zones (IZs) embedded in an otherwise homogeneous tissue (normal zone, NZ). Such as structure may be a cause of generating multiple re-entrant waves. The complexity of the IZs is due to electrophysiological parameter gradients within their spatial structure defining a central IZ (CIZ) surrounded by an ischemic border zone (IBZ). The wider organ level potentially random distribution of IZs further aggravates the heterogeneity and complexity [5, 6]. This preliminary study presents a relationship between the number of IZs and incidence of multiple re-entrant waves in a 2D model of the human ventricle.

Computational cardiology can provide mechanistic insights into the phenomenon of concern. It can provide a form of ground truth for clinical or experimental hypothesis, e.g. see [9, 10]. Increasingly, computational platforms are being designed to assist in clinical investigations [11, 12]. Mechanistic modelling of organs such as the heart draws upon cross disciplinary developments in computer science, biophysics, as well as applied mathematics, and is an ideal high performance computing (HPC) application.

## Methods

A 2D electrophysiological model was constructed to study the effects of post-dialysis ischemic heterogeneity. To do so, an existing human ventricle cell model was modified to permit simulation of the known effects of ischemia. The cell model was incorporated into a 2D model. Heterogeneity was incorporated into the 2D model by implementing ischemic zones, which permitted simulation of re-entrant wave breakup, or arrhythmia, under heterogeneous conditions.

### Clinical data in cell modelling

The 39 variable cell model of the human ventricle [13] was adopted to simulate cellular electrical excitations, called action potentials, under normal and ischemic conditions. The cell model electrophysiology was adapted by incorporating an additional adenosine triphosphate (ATP) sensitive potassium current, I_KATP_ [14]. The cell model was used to simulate normal and ischemic action potentials. To simulate the effect of ischemia, an increase of extracellular potassium, [K^+^]_o_, from 5.4 mM to 12 mM, an increase of I_KATP_ current conductance, g_KATP_, from 0.04 pA/pF to 2.05 pA/pF, and a 25% reduction of the upstroke current (sodium and calcium currents) conductances were implemented [7, 14–16]. The [K^+^]_o_ values were deduced from pre and post-dialysis serum measurements from 33 early dialysis patients (anonymised data), and adjusted to permit arrythmogenesis due to heterogeneity. Action potentials in NZ and CIZ are illustrated in Figure 1A.

**Figure 1.**
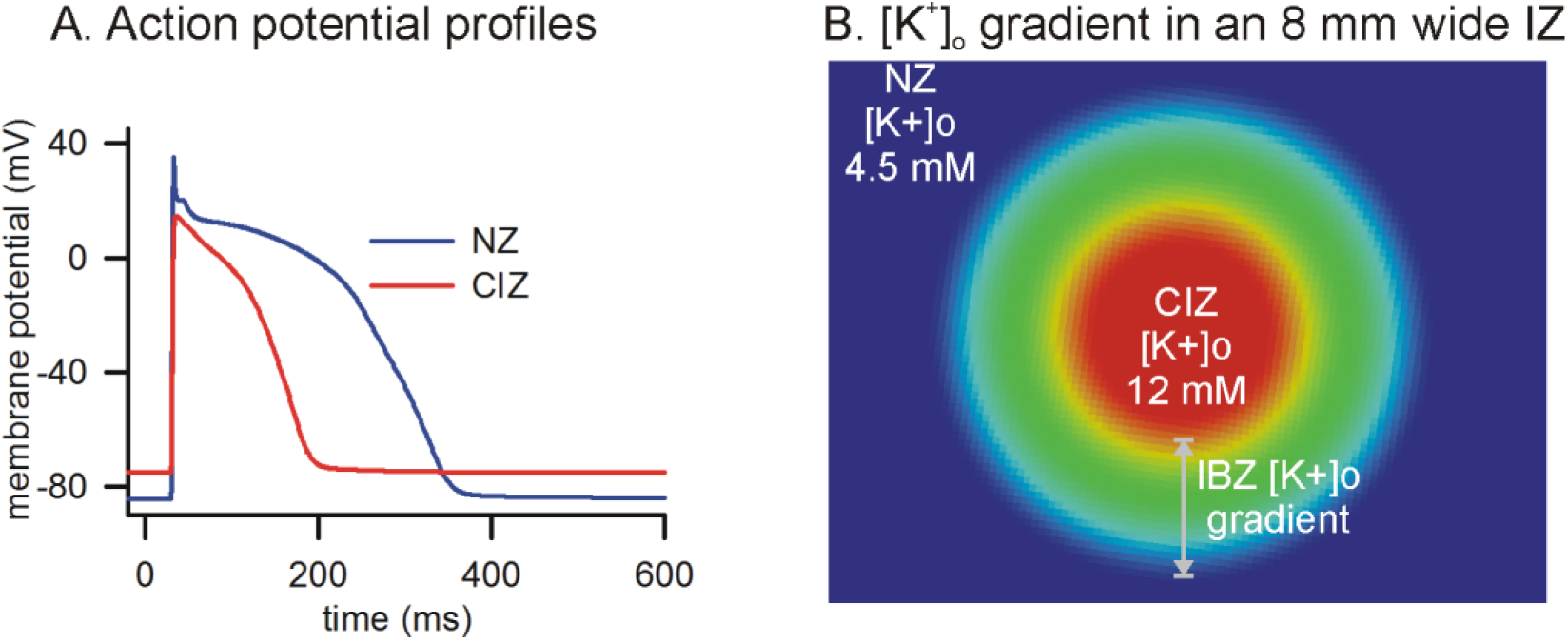
Simulated action potentials and IZ morphology. A: Action potentials in the NZ (blue line) and in the CIZ (red line). B: Potassium, [K^+^]_o_, gradient in one IZ of the 2D tissue. In NZ (blue tissue) [K^+^]_o_ was kept at 4.5 mM. In the CIZ (central red tissue), [K^+^]_o_ was kept at 12 mM. In the 2 mm wide IBZ, a linear gradient of [K^+^]_o_ was between 4.5 mM and 12 mM was implemented.

### 2D tissue model based on clinical imaging

Electrical wave propagation was simulated in a homogeneous 2D tissue model of the human ventricle. The model ventricle consisted of a structured finite difference grid spanning a 80 mm × 80 mm sheet (400 × 400 nodes, 0.2 mm internode distance). CT imaging data from an ongoing clinical trial (unpublished data) provided the dimensions and distribution of low perfusion IZ regions in the ventricle. The data was incorporated into the model as circular 8 mm wide IZs. Each IZ consisted of a CIZ surrounded by a 2 mm wide IBZ. The CIZ was assigned ischemic values as described above. Linear gradients of conductances and ATP where implemented in the IBZ between the CIZ and the NZ as described in the literature [8, 17]. The gradient of [K^+^]_o_ is used to illustrate the IZ consisting of CIZ and IBZ in Figure 1B.

The diffusion in the NZ was assigned a value of D = 0.004 mm^2^/ms to produce a conduction velocity of 0.5 m/s. Within the IZ, the diffusion parameter was reduced 10 fold representing a drastic reduction of ischemia induced cell-cell electrical coupling.

### Initial conditions and protocols to permit quantification of arrhythmia

The 2D geometry consisted of either a homogeneous NZ sheet, or a sheet with randomly distributed non-overlapping IZs. The random distribution of IZ in the 2D model is an approximation of our current observations in the imaging data. The geometry devoid of IZs was considered to be the pre-dialysis state of the ventricle. Arrhythmia is often quantified by studying a re-entrant wave’s tip trajectory, as well as the number of simultaneous re-entrant waves present in the ventricle. The post-dialysis geometries had 1, 10, or 20 IZs. To facilitate assessment of arrhythmia potential, i.e. genesis of multiple re-entrant waves, an initial single re-entrant wave (single phase singularity) using the phase distribution method as described previously [18, 19] was used to initiate a dynamical single re-entry. In brief, a phase, *ϑ*, between 0 and 2π was assigned to each location in the 2D sheet using the equation:

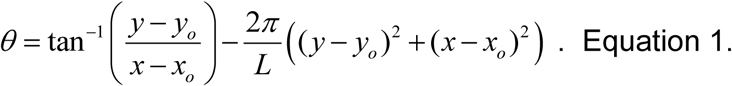

Where *L* is a distance measure in the re-entrant wave kept large for a single armed re-entry, and (*x_o_*, *y_o_*) is the user defined centre (phase singularity) of the re-entrant wave’s phase. In this study, L = 1000 mm and (*x_o_*, *y_o_*) = (40 mm, 40 mm). The NZ action potential in Figure 1, A was also assigned a phase value where *ϑ* = 0 corresponded to the value of all 39 dynamical variables in the cell model at t = 0, and *ϑ* = 360° corresponded to t = 600 ms. Using the spatial distribution obtained from Equation 1, the dynamical variables were distributed over the complete 2D sheet to initiate a single re-entrant wave. These initial conditions represent an artificially induced single tachycardia to assist with the study of genesis of multiple re-entrant waves. Whereas the physiological pacing rate of the human ventricle is around 1 Hz, re-entrant waves often pace the heart at a much higher pacing rate. Therefore, physiological pacing was considered to be overdrive suppressed by the induced propagations. The wave dynamics were permitted to evolve for a simulated time period of 2 seconds, with the membrane potential distribution in the 2D model being recorded at every 1 ms intervals.

### Pre and post-dialysis simulations

Simulations were performed under approximate pre-dialysis and post-dialysis conditions. In the pre-dialysis simulation, the 2D sheet was taken to be homogeneous without any IZs. In the post-dialysis simulations, four cases were considered where 1, 10, or 20 IZs were included into the 2D sheet. Within the confines of this study, the inclusion of an increasing number density of IZs may be interpreted as vascular disease progression.

### Quantification of arrhythmia

To quantify arrhythmia, two measures were applied. Firstly, the re-entrant wave tip trajectories were computed using a phase singularity detection algorithm [20] and developed in a previous study [21]. A re-entrant wave tip provides the location of a re-entry. Secondly, the dominant frequency (DF) at each location in the 2D model was computed. Dominant frequency was computed using algorithms available in the FFTW3 parallel library [22] permitting rapid as well as high throughput computation of dominant frequency maps. Due to the signal being composed of real numbers spanning the simulation, the discrete Fourier transform is computed more rapidly as well as using minimum required memory. This is due to the inherent Hermitian redundancy in the data.

### Algorithms and numerical methods

In this study, a mono-domain formulation with no-flux boundary conditions was adopted:

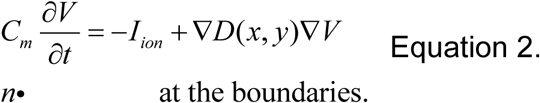

The overall wave propagation with the above equation was achieved using an operator splitting method. In this method and at each time step, the cell model is solved to obtain a partial solution. This partial solution is then used in the spatial diffusion part to obtain the full solution at the next time step.

Cardiac cell models consist of non-linear stiff ordinary differential equations (ODEs). The cell model equations were solved numerically using Backward Difference Formulae available in the Sundials library [23]. Electrical wave propagation was simulated in the spatially extended models using reaction-diffusion partial differential equations (PDEs). The PDE was solved using second order explicit finite difference method, encoded into a message passing interface (MPI). The spatial solver exploits functions from the Portable, Extensible Toolkit for Scientific Computation (PetSc) library, that permitted accurate and stable solutions based on Newton iterations [24, 25]. Whereas the ODE as well as the PDE solvers both use variable time steps to attain user defined tolerances, the maximum time step was set to 0.1 ms. The parallel code is implemented in the C language using the message passing interface (MPI) to enable use of robust scientific libraries, be extensible, and to exploit MPI parallelisation. The code was profiled using Allinea MAP software available in Compute Canada HPC supercomputers.

## Results

The cause-effect relationship was established by performing pre-dialysis and post-dialysis simulations. The simulations permitted the quantification of re-entrant wave breakup in terms of number of re-entry tips and alteration of dominant frequency.

### Homogeneous pre-dialysis ventricle does not promote arrhythmia or multiple re-entrant waves

The pre-dialysis case is illustrated in Figure 2. The re-entrant wave meander was found to be limited, due to the overall slow conduction velocity in the tissue. The absence of IZ heterogeneities in the 2D sheet resulted in a rotating wave along its natural meander path. The re-entrant wave tip trace (Figure 2, B) shows that the meander area was small. Whereas such a tachycardia may self-terminate in a healthy ventricle, the persistence of this re-entry was due to the slow conduction. Further, the computed dominant frequency (Figure 2, C) shows that most of the 2D sheet was paced at the period of the rotating wave of around 3 Hz. A small region was paced at a lower pacing rate of around 2.5 Hz. This small region is enclosed by the meander pattern where the arm of the re-entry does not always reach.

**Figure 2.**
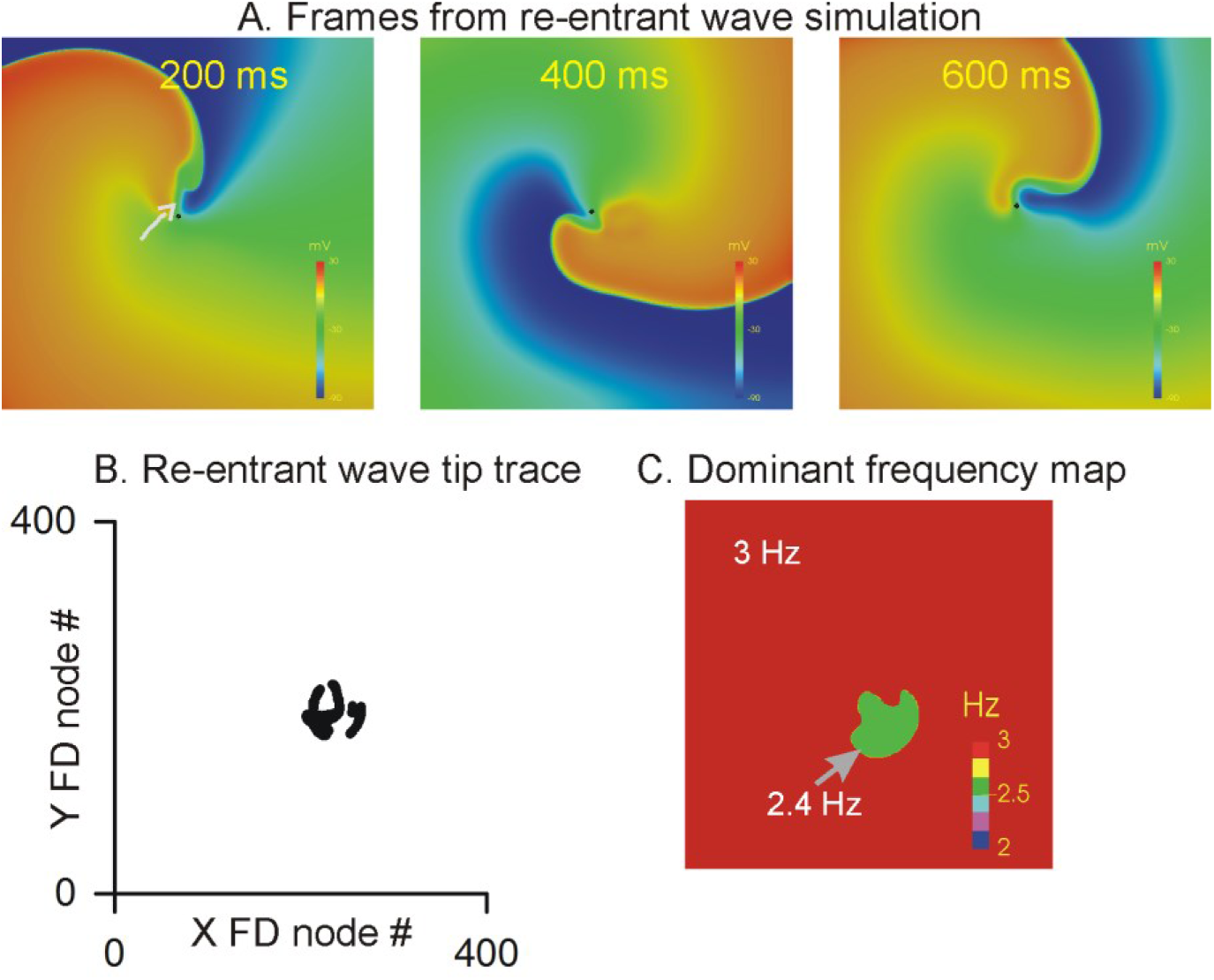
Pre-dialysis simulation. A: Representative frames from the 2 second simulation of a single re-entrant wave. The small dots show location of the estimated re-entrant wave tip. The colorbar represents a membrane potential range of −90 mV to +20 mV. B: Re-entrant wave tip trace throughout the span of the simulation. C: Dominant frequency map of the 2 seconds of electrical activity.

### IZ heterogeneity is a cause of arrhythmia by promoting re-entrant wave breakup

The results of post-dialysis simulations are illustrated in Figure 3. Increasing conduction heterogeneity was simulated by including 1, 10, or 20 IZs (Figure 3, A, top panels). Each IZ was constructed as described in the methods and Figure 1. The dominant frequency maps (Figure 3, A, middle panels) show that when the number of IZs was small (i.e. 1), the maximum pacing experienced by the tissue was around 2 Hz. However in the case when a significant portion of the tissue included IZs, i.e. 20 IZs, the maximum frequency was significantly higher at 6.5 Hz. The fast rate of pacing occurred exclusively in the IBZs as shown in Figure 3, A, middle panels. Further, the tip traces of the re-entrant waves show that in addition to the induced single tachycardia, the IBZ always promoted short lived but many re-entrant waves.

**Figure 3.**
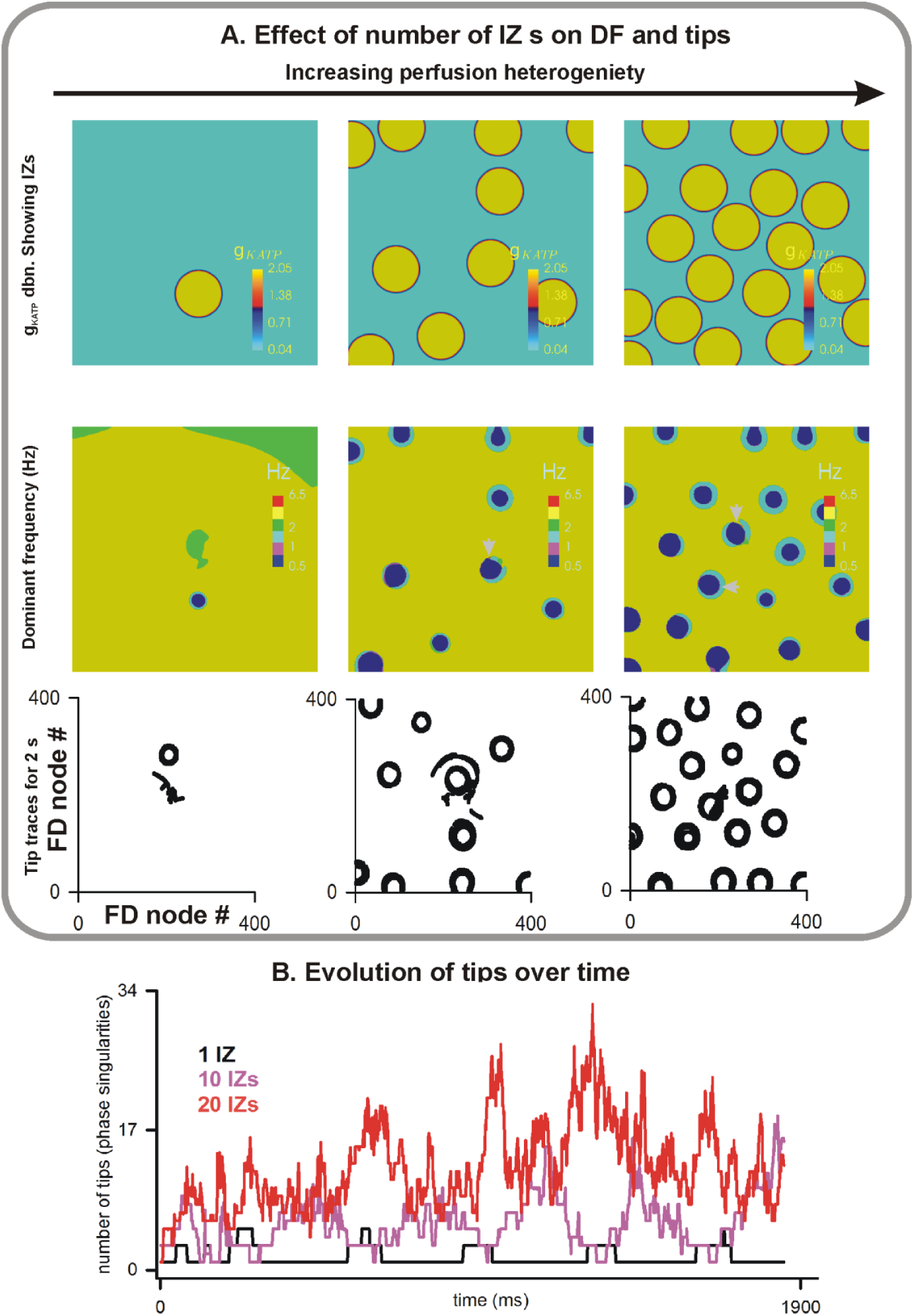
Post-dialysis simulation under various levels of ischemic heterogeneity. A: Top panels show the distribution of the IZ patches. The gradient of the I_KATP_ conductance, g_KATP_, is shown in the individual panels. Middle panels show the corresponding dominant frequency maps. The colour bar represents frequencies between 2 Hz and 7 Hz. Bottom panels show the re-entrant wave tip patters. B: Number of re-entrant wave tips as a function of time in the 3 cases shown.

This number of re-entrant wave tips as a function of time are summarised in Figure 3, A, bottom panels and shown explicitly in Figure 3, B. Further, Figure 3, B shows that the number of re-entry tips oscillated between a maximum and minimum in case of 1 or 10 IZs. This indicates that the induced single tachycardia was the dominant rotor driving the generation of smaller phase singularities in the IBZs. However, when the number of IZs was large, the induced tachycardia rapidly dissipated giving rise to erratic wave propagations dictated by the IZ locations and their IBZs. Although the number of tips was large in the 20 IZs case, the time evolution still shows an oscillation which is due to the initial conditions affecting the short 2 s simulation. The maximum number of re-entrant wave tips induced by one IZ was around 10. However, the maximum number of re-entrant wave tips induced by 10 IZ patches was 20, while in case of 20 IZs it was 34.

The model code took approximately 11 hours using 24 processors to simulate one case, and required 0.36 GB memory. To identify the bottlenecks, the most important functions of the code were profiled using the MAP profiler (Figure 4). At each iteration, it was found that the maximum CPU time was spent during the solutions of the ODE and PDE parts of the model. In both solvers, ODE and PDE, a numerical approximation of the relevant sub-systems Jacobian is computed automatically using the respective library functions. This process consumed 70% of the total time/effort. The computation of the Jacobian provides the mechanism for accurate Newton iteration based solutions to defined tolerance values. This results in the actual calculation of solution at the next time step to be 23% of total CPU effort. In previous codes a major MPI bottleneck was the collection of output data to the master rank, and the master rank outputting the data while all other processes were held. In our code due to parallel output, the I/O overheads were found to be negligible.

**Figure 4.**
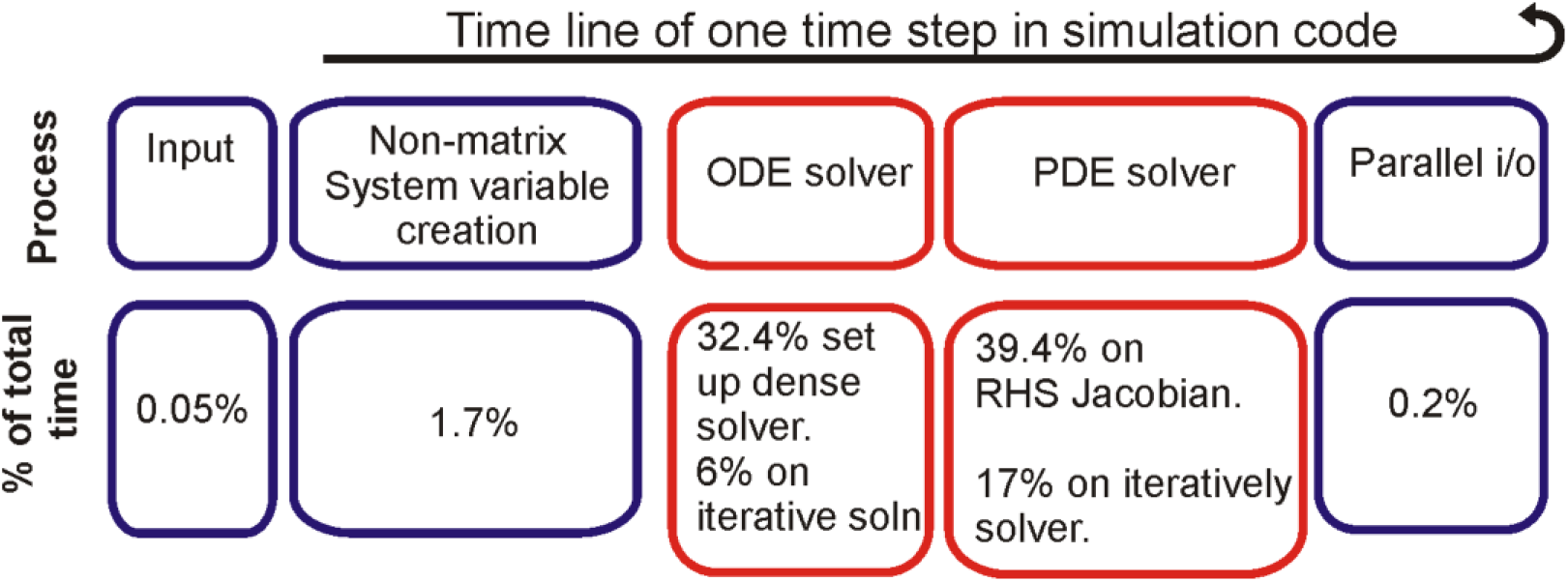
Efficacy profile of the code. Top panels show the main components of the MPI based code developed in our group. Bottom panels show the corresponding time taken by each component. (see text for discussion).

## Future directions

### Modelling limitations

As a first study, we have used simple 2D sheet model. Combined with the biophysically detailed cell model, it permitted an exploration of the arrhythmia potential in the human ventricle.

The cell model was used to produce normal and IZ action potentials. However, it is known that the human heart is transmurally heterogeneous in electrophysiological properties having epicardial (epi), mid-myocardial (mid), and endocardial (endo) action potentials (Figure 5, A) [26]. At the spatial level, the model geometry used in this study represents the ventricle as a 2D sheet. However, the 3D heart has an uneven geometry and includes several heterogeneity inducing features such as fibre orientation, Purkinje stimulation network, papillary muscles, and vasculature [27]. The structural heterogeneity becomes more relevant as we move towards computational study of cardio electro-mechanics [27]. The cardiac conduction system’s end points, called Purkinje insertion points, have significantly distinct electrical properties which increases the electrical heterogeneity (Figure 5, A). Further, the 0.1 mm long cardiomyocytes are orientation in a complex manner as quantified in extensive high resolution imaging studies [28, 29]. The transmural cellular organisation gives rise to an anisotropic diffusion regulated by the fire directions (Figure 5) which requires significant modification of Equation 2. Ischemia is a multi-factorial condition that requires inclusion of several cellular and sub-cellular functional alterations for an accurate health status assessment. In future studies, the sensibility and suitability of the models, as done in case of heart cell models [30–32], will be carried out prior to their use in mechanistic simulations. Ischemia is also known to affect the cardiac structure and micro-architecture [33], which are topics of future investigation in light of our clinical observations.

**Figure 5.**
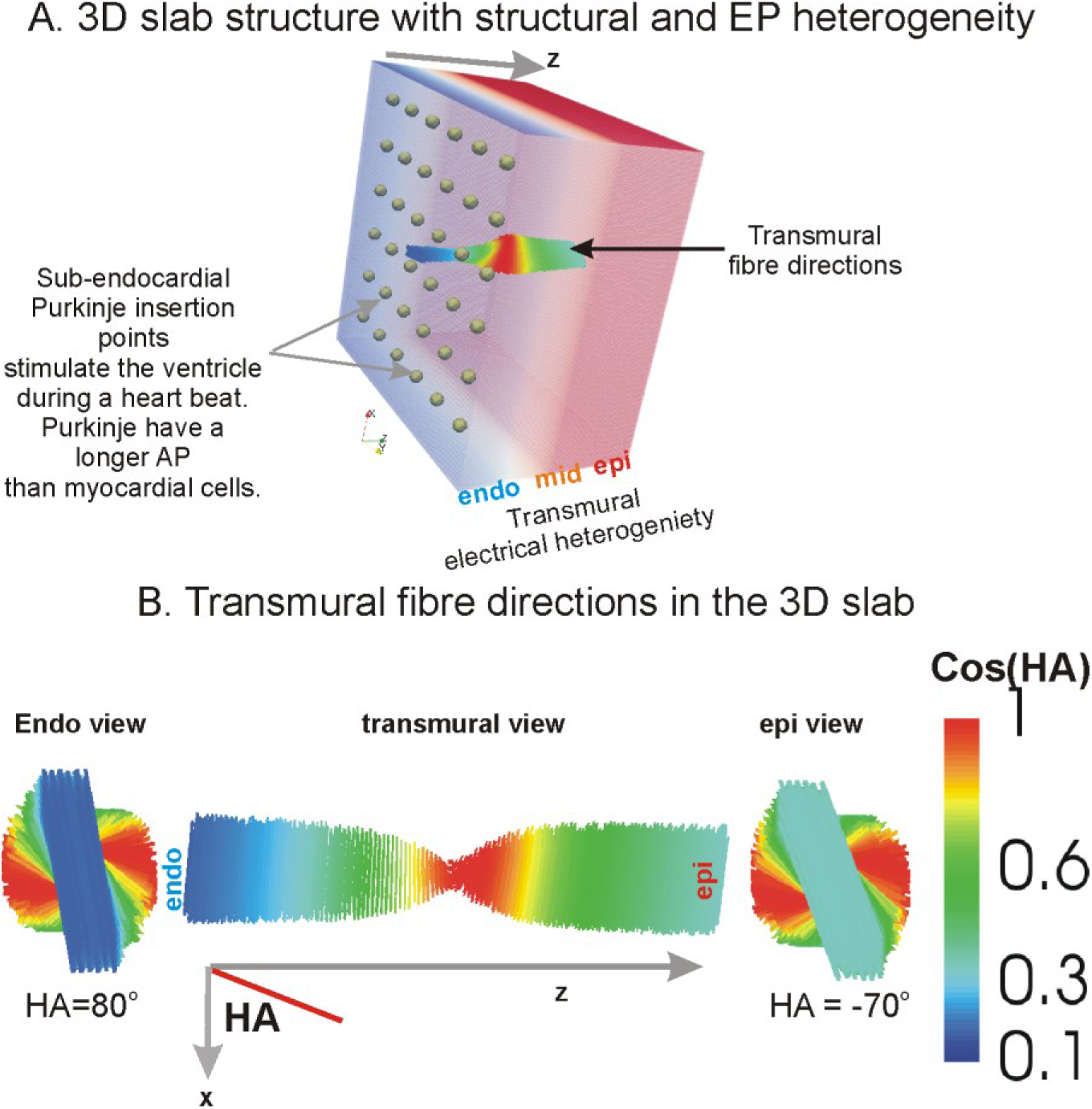
Human ventricle slab illustrating electrical heterogeneity and spatial anisotropy. A: The Purkinje end points (spheres separated by 3 mm) are located in the sub-endocardium. The transmural ventricular wall is composed of endocardial tissue type (endo), mid-myocardial tissue type (mid), and an outer epicardial tissue type (epi). The alteration of cardiomyocyte orientation is shown in the colour coded transmural fibre directions. B: Endocardial (from inside the ventricle cavity) view of the fibre directions is shown in the left panel. The middle panel shows transmural view, and the right panel shows the epicardial view. The fibres are colour coded according to the helix angle (HA) or the fibre vectors X component. In case of a rectangular slab, HA is the angle of the fibre with respect to a reference X axis.

### Computational aspects

A spectrum of computing developments is planned to permit real time patient specific assessments. In this study, the rectangular geometries that were used lend themselves to a straightforward box partitioning. It is known that realistic geometries occupy less than 30% of the total volume [34], which often leads to uneven load on processors in distributed memory machines. In addition, a larger amount of memory is required for full 3D simulations. To optimise memory use, a method for mapping of the dynamical system variables onto a 1D array was developed previously [34]. To further improve both the load balancing as well as memory use, graph theory based partitioning will be implemented in the future [35]. With significance to box geometry based studies such as the present, it is relevant to provide user defined functions rather than relying on their numerical computation to improve efficiency of the codes. The accuracy may also be improved in the case when the Jacobians provide closed form equations for the relevant derivatives.

## Conclusions and Discussion

The main findings of this study were: 1) The electrophysiological heterogeneity in the IBZs causes generation of re-entrant waves; 2) the number of IZs is directly proportional to both measures of arrhythmia potential, i.e. number of re-entrant wave tips and dominant frequency.

This study presents a relationship between IZs and degeneration of a single tachycardia into ventricular fibrillation. In pre-dialysis patients’ hearts, the myocardium may be diseased but may also be homogeneous because the serum has the opportunity to diffuse everywhere. On the other hand, post-dialysis serum content is likely to be significantly altered due to the 4 hour long dialysis induced ischemic shock. The post-dialysis refreshed serum is unlikely to permeate the complete myocardium due to a combination of vascular disease and other unidentified loss of function. In particular, our study shows that significant fibrillation occurs when the number of IZ is large (Figure 3). This finding is in line with other theoretical studies that highlighted the role of tissue spanning fibrosis [5, 6], which is another cause for conduction heterogeneity.

Whereas a reasonable number of simulations were performed in this study, the number of modelling parameters affected by myocardial disease is large. For instance, a study by Kharche et al. explored the spatial meander of tightly rotating scroll waves (3D re-entrant scroll waves) in the 3D atrium (Figure 6) [18]. We showed that the scroll waves were affected by the atrial geometry’s thickness and unevenness. Each scroll wave trajectory shown in Figure 5 required the use of 144 processors for over 48 hours. It can be appreciated that real time detailed simulations may become possible when issues such as identified in Figure 4, as well as other hitherto unidentified issues are addressed. Further to the algorithmic developments, it is also necessary to exploit the upcoming GPU and GPU-CPU hybrid technologies to accelerate the simulations [36, 37].

**Figure 6.**
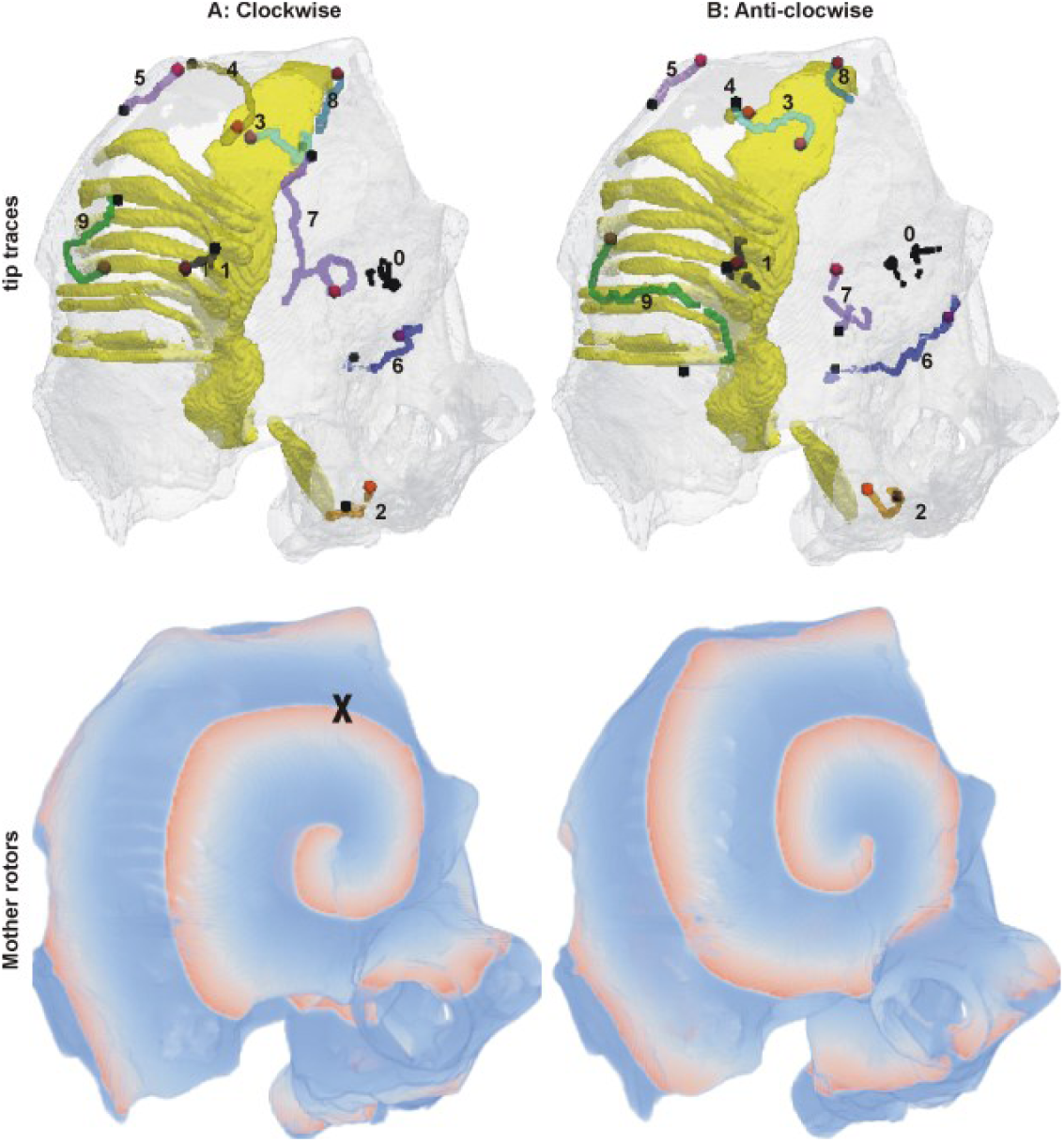
Meander of tightly rotating scrolls due to uneven 3D heart anatomy. Exemplar large scale simulation of pathological electrical activity in the human atrium (upper chambers of the heart), please see [18] for details (reproduced under Creative Commons Attribution License). Top panels: Black dots denote initial locations of scroll waves, whereas red dots denote final locations after a simulated time of 40 seconds. The lines joining the start and end points show the trajectory taken by the scroll wave during the 40 s. Bottom panels show one representative frame from the clockwise (left) and anticlockwise (right) simulations.

Cardiac ischemia is a hotly studied clinical condition using clinical, experimental, and computational approaches [17]. The ongoing basic science efforts in our clinical-computational group reflect the necessity to advance our knowledge of this widespread clinical condition that commonly occurs in our hospitals. It is evident that extension of the existing technology to benefit the clinical sciences requires both, intense modelling and specific experimental and imaging input to personalise the outcomes. Mechanistic computational modelling is expected to play a role within health sciences research and practice as summarised in Figure 7, which calls for extensive cross-disciplinary collaboration, including that with HPC specialists.

**Figure 7.**
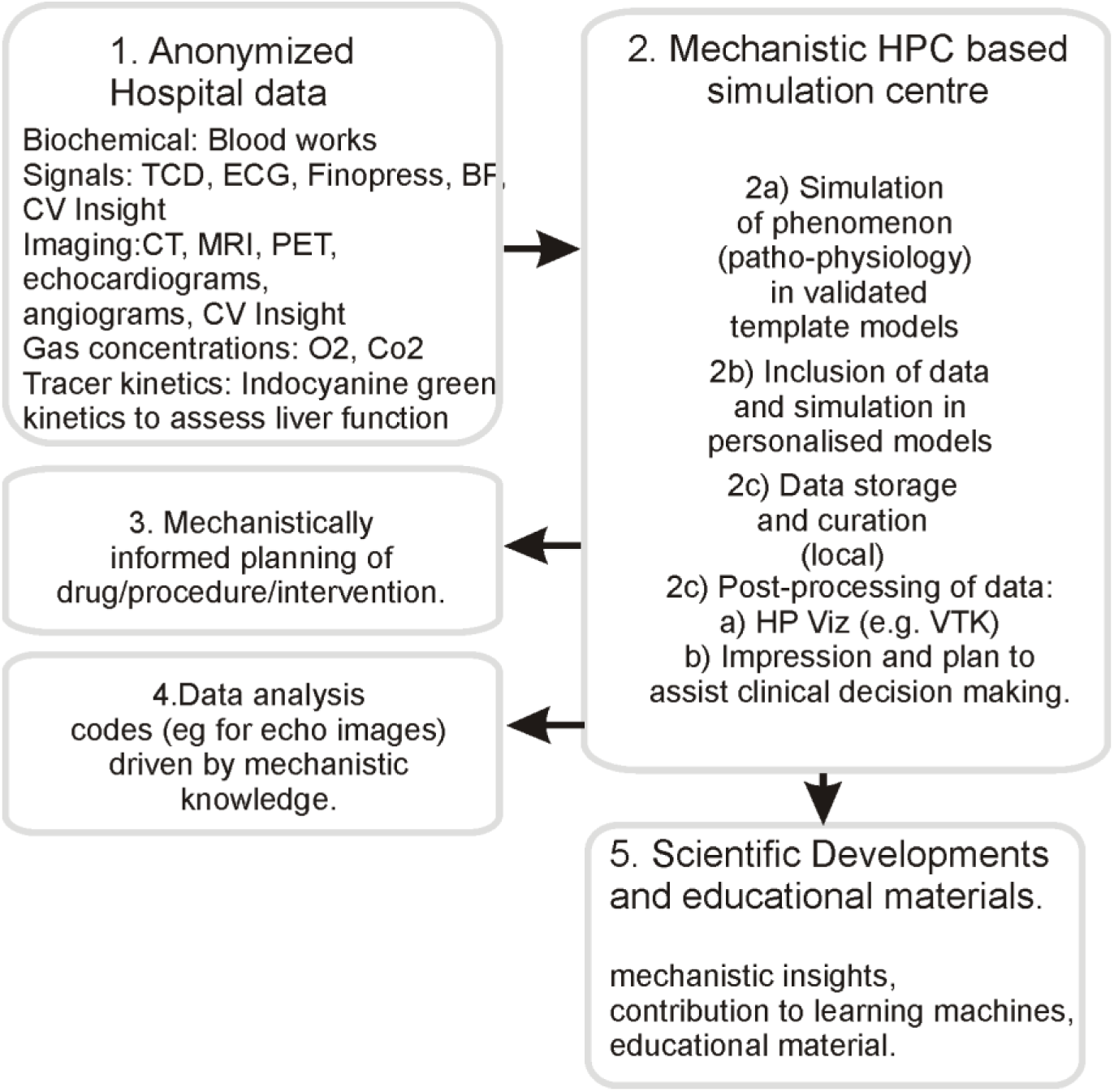
Proposed work flow and potential outcomes in HPC based clinical decision making.

## Author contributions

SRK and CWM designed the study. In consultation with CWM, SRK implemented the simulations. SRK performed the simulations, data analysis, and code profiling. SRK wrote the draft, and SRK and CWM finalised the manuscript after an internal review.

## Acknowledgements

SRK is supported by a grant held by CWM. CWM holds multiple grants for clinical and basic science research. We thank LHSC IT Services for their support in installing hardware. We thank SHARCNET for HPC resources to SRK. Ms Tanya Tamasi obtained the anonymised serum data from the LHSC database. We thank Drs. Elena Qirjazi and Marat Slessarev for discussions and critical comments.

